## Supplementary Figure 1 for "PySTACHIO: Python Single-molecule TrAcking stoiCHiometry Intensity and simulatiOn, a flexible, extensible, beginner-friendly and optimized program for analysis of single-molecule microscopy data"

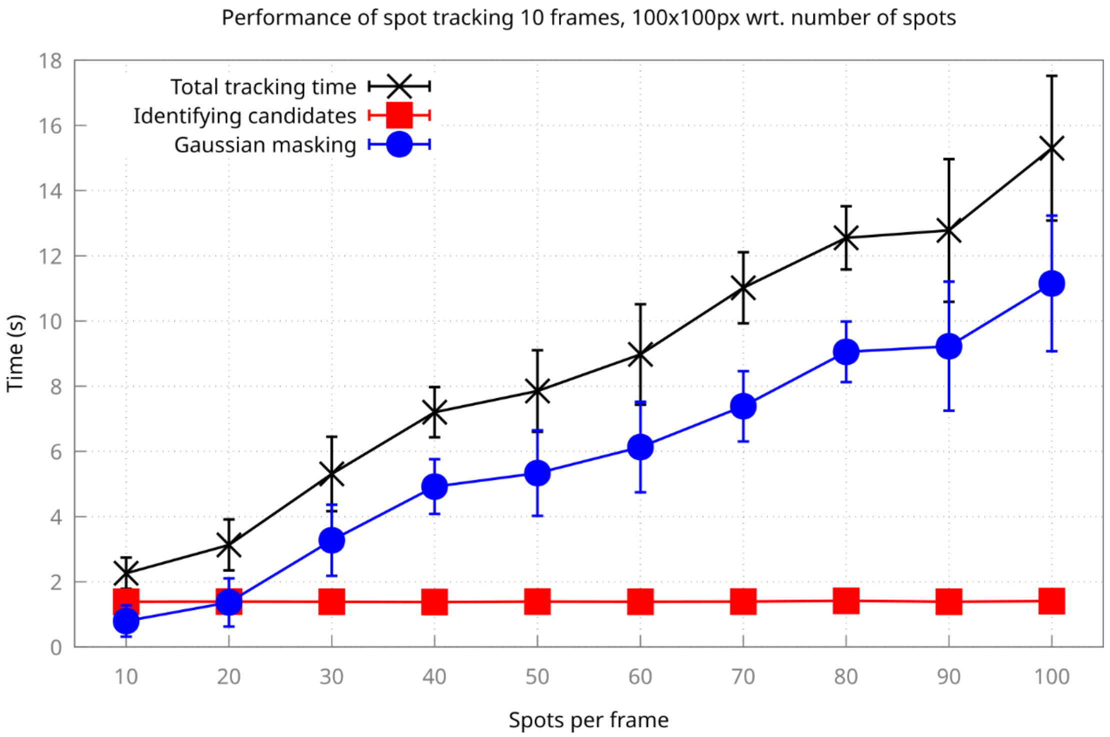


Supplementary Figure 1: Performance of PySTACHIO with increasing spot density (where a "spot" is tracked simulated diffraction-limited fluorescent focus). A simulated frame of 100x100 pixels was simulated with varying numbers of spots. PySTACHIO's performance overall remains almost linear with the majority of the run time being taken up with the iterative Gaussian masking routine.
